# Improved protein complex prediction with AlphaFold-multimer by denoising the MSA profile

**DOI:** 10.1101/2023.07.04.547638

**Authors:** Patrick Bryant, Frank Noé

## Abstract

Structure prediction of protein complexes has improved significantly with AlphaFold2 and AlphaFold-multimer (AFM), but only 60% of dimers are accurately predicted. A way to improve the predictions is to inject noise to generate more diverse predictions. However, thousands of predictions are needed to obtain a few that are accurate in difficult cases. Here, we learn a bias to the MSA representation that improves the predictions by performing gradient descent through the AFM network. We effectively denoise the MSA profile, similar to how a blurry image would be sharpened. We demonstrate the performance on seven difficult targets from CASP15 and increase the average MMscore to 0.76 compared to 0.63 with AFM. We evaluate the procedure on 334 protein complexes where AFM fails and demonstrate an increased success rate (MMscore>0.75) of 8% on these hard targets. Our protocol, AFProfile, provides a way to direct predictions towards a defined target function guided by the MSA. We expect gradient descent over the MSA to be useful for different tasks, such as generating alternative conformations. AFProfile is freely available at: https://github.com/patrickbryant1/AFProfile

## Introduction

Protein structure prediction of single chains is now highly accurate with AlphaFold2 (AF) [1]. The prediction of multimeric protein structures is not, as recent benchmarks report an average success rate (MMscore [2] above 0.75) of around 60% for dimers and the accuracy decreases with the number or chains [3]. By adapting the information of the multiple sequence alignment (MSA) that goes into AF, the network which is trained only on single chains can be adapted to outperform all other protein docking methods [4–6]. AlphaFold-multimer (AFM) uses the same adaptation of information but is trained in an end to end fashion obtaining slightly better performance [7].

Recycling significantly improves performance for both single- and multi-chain proteins [1,4]. In each recycling iteration, the input MSA is sampled to create input features (as the whole MSA can’t fit inside the network), embedding more information inside the network [7]. By making many predictions (6000) and adding noise, the accuracy can go from poor to almost perfect in some cases[8,9]. Further, the confidence metrics provided through the predictions can separate the accurate cases from the inaccurate ones [10]. Together, this suggests that:

1. The information to make high quality predictions is available in the MSAs.
2. The algorithm of AFM is sufficient to generate highly accurate structures given the correct information.
3. Accurate structures can be selected through quality metrics.

The main problem is therefore to find the right MSA information so that AFM can utilise it to generate high-quality predictions and further algorithmic advances may not be necessary.

The issue of how to find a combination of sequences with sufficient information to predict an accurate structure is difficult. During recycling, a combination of different samples of sequences taken from the MSA is built up. The cumulative representation resulting from this sampling, including the internal representation kept through the recycling, is what determines the outcome. As this is a complicated collection of information, it is unclear how to determine an optimal selection beforehand, which is why sampling is performed randomly [1,7].

A method to enhance the MSA information for the AFM network would potentially improve the accuracy of the protein complex predictions. As the quality metrics predicted by the network serve as a guide for model quality, we propose to use these to guide the representation of the MSA in the network to be more favourable. This idea is similar to advances in protein design which search for sequences that generate high confidence metrics predicted by AF [11–14]. The most efficient way to search for such a representation should be gradient based, i.e. by backpropagating through AFM. By doing that, we effectively denoise the MSA profile, similar to how a blurry image would be sharpened to become more clear.

## Results

### Improving the MSA profile with gradient descent

Structure prediction depends on coevolutionary information. Obtaining more ordered information greatly improves the performance of protein complex prediction [4]. The most important information that goes into AlphaFold-multimer (AFM) is the multiple sequence alignment (MSA) cluster profile. This is a statistical representation of which amino acids are likely to be found at certain positions in an MSA. The importance of the profile can be shown by setting a random MSA profile and keeping all other MSA features intact. The result is a completely disordered structure. If a structure can be destroyed by corrupting the profile, perhaps it can be improved by finding the most useful features in the evolutionary representation, i.e. *denoising it*.

In CASP15, the highest performing group (Wallner [8]) achieved their performance by introducing noise into AFM and making many predictions (6000 in total) [8,15]. Although high performance was achieved in many cases, this procedure can not be deployed at scale due to the high computational requirement. Moreover, there is no guarantee that a good model will be the outcome as sampling a good MSA can be a rare event. To solve these issues we created a way to improve the MSA profile to obtain more accurate structures, a process which we call **AFProfile** (Figure 1a).

**Figure 1.**
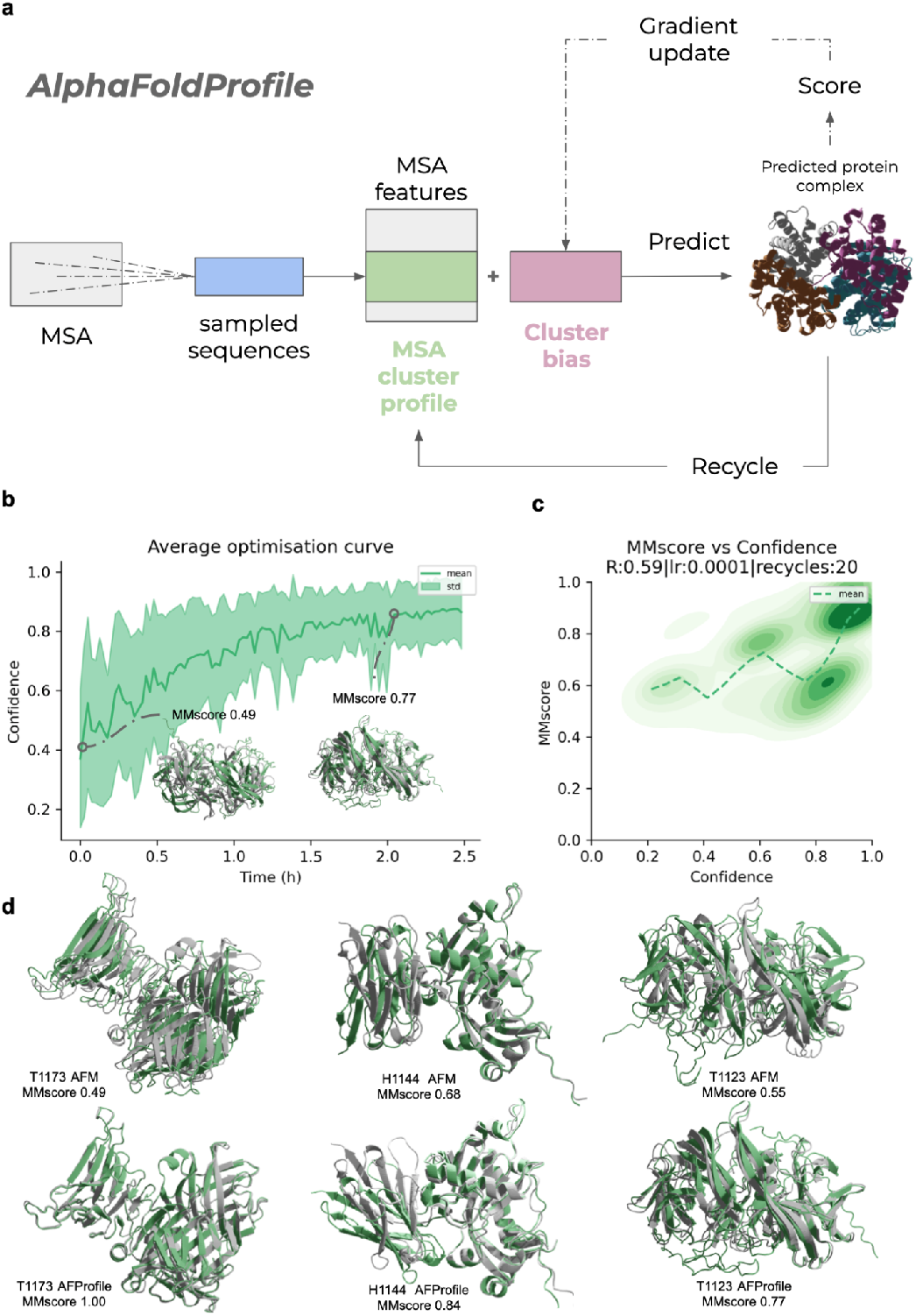
a) The AFProfile Method. Starting with MSAs generated by the default AlphaFold-multimer pipeline, sequences are sampled and MSA features are created. These features are used to predict the structure with AlphaFold-multimer and among the most important is the cluster profile. We learn a residual to this, the cluster bias, which effectively denoises the MSA profile into a representation that generates a higher confidence score and a more accurate structure. Recycling operations are performed to embed more information in the network and the cluster bias acts across the recycles. This can be seen as denoising a blurry image to make it sharper. **b)** Average predicted model confidence vs time in hours on one NVIDIA A100 GPU using gradient descent for the 7 CASP15 targets (H1134, T1123, T1173, H1141, H1144, H1140 and T1187). A learning rate (lr) of 1e-4 with the Adam optimiser and 20 recycles were used here (Methods). Example models for T1123 (green) are shown at different points of predicted confidence with the top ranked CASP15 model in structural superposition (gray). In total, 100 optimisation steps were performed per target (n=700). **c)** Confidence vs MMscore across the 7 CASP15 targets (H1134, T1123, T1173, H1141, H1144, H1140 and T1187). For each target, 100 optimisation steps were performed (n=700). The Spearman R is 0.68, the lr 1e-4 and the number of recycles used set to 10 (Methods). A density plot using all samples (n=700) and a running mean using a step of 0.05 confidence are shown. **d)** Examples from CASP15 with the best prediction in grey and AlphaFold-multimer(AFM) and AFProfile coloured green for targets H1144, T1123 and T1173. For T1173, the MMscore improves from 0.49 with AFM to perfect with AFProfile (1.0). For H1144, one of the chains is in the wrong orientation, while the right configuration is found with AFProfile (MMscore=0.84). For T1123, both chains are slightly wrong (MMscore=0.55), while AFProfile improves the score to 0.77 (accurate model >0.75).

AFProfile adds a bias to the MSA profile which acts as a correction term. The bias is created by performing gradient descent through the AFM network with the objective to maximise the confidence (equation 1) AFM has in the predicted structure. The confidence that AFM predicts has been shown to correlate with model quality [3,8]. The process can be seen as denoising the MSA representation, similar to how a blurry image would be sharpened.

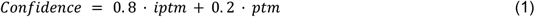

Here, iptm is the predicted TM-score [16] in the interface and ptm that of the entire complex.

We optimised the procedure on seven targets from CASP15 in which the Wallner group outperformed the standard AFM prediction using AFsample (Methods). The bias addition being learned rapidly improves the confidence (Figure 1b) and we find that the confidence and MMscore trajectories correlate with a SpearmanR of 0.59 and a high density of structures with MMscores and confidences >0.8 (Figure 1c). AFM is outperformed with an average MMscore of 0.63 vs 0.76 for AFProfile (Table 1, Figure 1c). For three targets (H1144, T1123 and T1173) the models go from unsuccessful with AFM to successful (MMscore>0.75) with AFProfile (Figure 1d).

**Table 1.** Prediction results on 7 targets from CASP15 from the Wallner group, AFM and AFProfile. The AFsample MMscores are higher on average, 0.97 vs 0.76 for AFProfile. Compared to AFM, the increase in MMscore is 0.13 on average (0.76 vs 0.64), making 3 additional targets successful (MMscore>0.75). The successful models (MMscore>0.75) are marked in bold.

| Target | Reference MMscore (top model from CASP15) | AFsample MMscore | AF-multimer MMscore | AFProfile MMscore |
| --- | --- | --- | --- | --- |
| H1134 | **0.97** | **0.97** | **0.92** | **0.88** |
| H1140 | **0.97** | **0.97** | 0.62 | 0.61 |
| H1141 | **0.98** | **0.97** | 0.68 | 0.68 |
| H1144 | **0.99** | **0.99** | 0.68 | **0.84** |
| T1123 | **0.89** | **0.91** | 0.55 | **0.77** |
| T1173 | **0.99** | **0.99** | 0.49 | **1.00** |
| T1187 | **0.98** | **0.96** | 0.49 | 0.51 |
| Average | **0.97** | **0.97** | 0.63 | **0.76** |

Compared to using AFsample, the performance of AFProfile is lower (average MMscores of 0.76 vs 0.97). Learning the residuals to the cluster bias speeds up the prediction at least 60-fold on average as 100 iterations are used compared to 6000 with AFsample. Since AFM v2.3 uses 100 models with 20 recycles [17] this process proves as efficient as the default AFM. Most importantly, AFProfile provides a way to direct the predictions towards more accurate complexes, and potentially alternative conformations (see below).

### Improving the success rate of AlphaFold-multimer on difficult targets

As we developed the learning of the MSA cluster bias on only 7 targets from CASP15, we now set out to evaluate the real expected performance of AFProfile on a much larger set of 334 nonredundant complexes with 2-6 chains not included in the training set of AFM [7]. These were selected from a recent benchmark of complexes as AFM fails to predict them with sufficient accuracy (MMscore<0.75) [3]. The success rate (MMscore>0.75) increases to 8% (26 complexes) for these difficult targets with AFProfile (Figure 2a).

**Figure 2.**
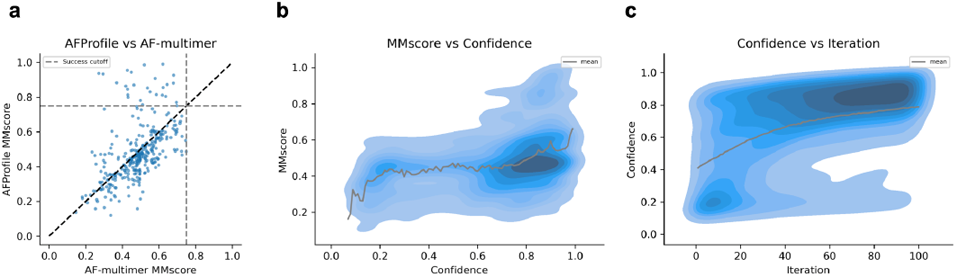
**a)** MMscores of difficult targets that fail (n=334) with AF-multimer compared to AFProfile. In total, 7.8% of the failed examples can be rescued (MMscore>0.75) with AFProfile. **b)** MMscore vs confidence. The density represents all predictions (100 per target, n=33400) and the line the running mean using a step size of 0.01 confidence. **c)** Density plot of the confidence vs iteration of gradient descent with AFProfile (n=33400). At higher iterations, there is a strong density of high confidences. Following Figure 2b, this region is more likely to have high MMscores.

A similar relationship between the MMscore and confidence as for the CASP15 set is observed. For confidences below 0.8, the average MMscore is 0.4 and only increases to 0.6 for confidences around 0.9. However, most complexes are not predicted accurately here resulting in a concentration of complexes with confidences of 0.8-0.9 and MMscores of 0.5, suggesting a higher difficulty in this set (Figure 2b). Figure 2c displays the relationship between the confidence and the iteration (gradient descent step). At higher iterations, there is a strong density of high confidences. Following Figure 2b, this region is more likely to have high MMscores. This suggests that the optimisation procedure does improve the outcome and that running more iterations is favourable.

#### Structures with larger changes are more successful

As most complexes reached a confidence of >0.8, but their MMscores did not improve to above 0.75 (Figure 2a), it is possible that AFM will not be able to predict these at all (i.e. these targets are too difficult). It is also possible that an alternative conformation is predicted that is currently unknown but this can not be verified. Figure 3a shows the final MMscore vs the structural change during the optimisation procedure with AFprofile as measured by the change in MMscore (ΔMMscore). In most cases, the change in MMscore is small. Even in cases where this change is small, it is important to select correct models from the confidence scores. It is unlikely to observe a substantial decrease in the MMscore with the optimisation procedure suggesting that when a high MMscore is obtained initially, a similar prediction is maintained.

**Figure 3.**
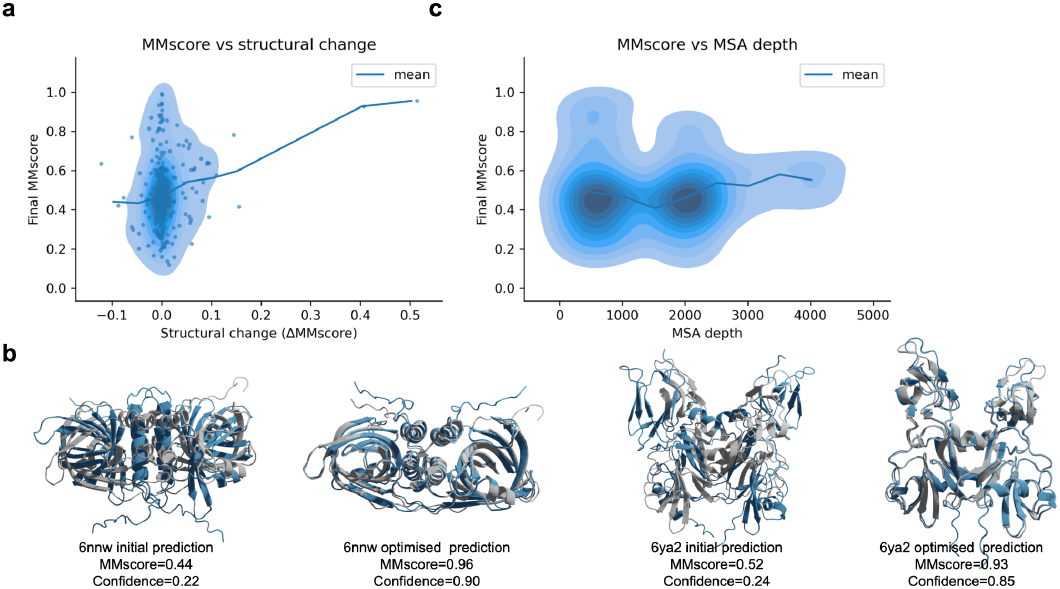
**a)** Final MMscore vs structural change during the optimisation procedure as measured by the change in MMscore (ΔMMscore). The density and points represent all predictions (n=334) and the line the running mean with a step size of 0.05 ΔMMscore. This shows that the MMscore is unlikely to decrease with the optimisation procedure and that when the structural change is large compared to the initial prediction (measured by ΔMMscore), it is likely to obtain a higher MMscore. This is expected as the predicted structures are not accurate using AFM (MMscore<0.75). **b)** Initial and AFProfile optimised structure of PDBIDs 6nnw (https://www.rcsb.org/structure/6NNW) and 6ya2 (https://www.rcsb.org/structure/6ya2). The native structure is in grey and the predicted ones in blue. The MMscores increased from 0.44→0.96 and 0.52→0.93 and the confidences from 0.22→0.90 and 0.24→0.85, respectively, during the optimisation. **c)** Number of sequences in the MSA input representation generated by the AFM pipeline (MSA depth) vs the final MMscore after optimisation with AFprofile. The density represents all predictions (n=334) and the line the running mean with a step size of 500 in MSA depth.

When the structural change is large compared to the initial prediction it is more likely to obtain a higher MMscore. This suggests that the initial conformations generated by AFM are wrong in these cases and that AFProfile recovers other conformations that are more likely to be correct. The relationship between the structural change and MMscore is expected as the predicted structures are not accurate using AFM (MMscore<0.75). Examples for PDBIDs 6nnw (https://www.rcsb.org/structure/6NNW) and 6ya2 (https://www.rcsb.org/structure/6ya2) are shown in Figure 3b.

In addition, we analysed the impact of the MSA depth (number of sequences in the input MSA representation from AFM) on the outcome (Figure 3c). It does not seem favourable to have any typical MSA depth. Likely, the quality of the information contained inside the MSA and not its size determines the outcome. This may explain why these structures can’t be predicted with sufficient accuracy as well. It may be that the MSAs do not contain sufficient information and, therefore, there is nothing for AFProfile to improve upon.

## Conclusions

AFProfile can enhance the outcome of structure prediction with AlphaFold-multimer (AFM) by denoising the MSA profile. The predicted confidence metric from AFM acts as a guide for this process and correlates with the MMscore of the complexes (SpearmanR=0.59 for the 7 CASP15 targets, Figure 1). Applying AFP to a large test set of 334 complexes where AFM fails, results in accurate predictions (MMscore>0.75) for 8% of the complexes. In some cases, it seems likely that these are alternative conformations to those generated by AFM as there is a relationship between the structural change and MMscore (Figure 3).

AFProfile, provides a way to direct predictions towards a defined target function guided by the MSA. We expect the procedure of using gradient descent over the MSA to be applicable to a variety of tasks such as generating alternative conformations. When the highest confidence prediction is searched, different structures are explored. This can be likened to traversing the loss landscape to find low energy valleys of putative alternative conformations [10,18,19]. This opens up the possibility to obtain conformational ensembles and to efficiently sample the dynamic landscape of alternative protein conformations for both single chain and multimeric structures.

## Methods

### Data

#### CASP15

To see if we can improve upon AlphaFold-multimer (AFM) and AFsample, we select eight targets from CASP15 where AFsample significantly outperforms AFM. These are H1134, T1123o, H1129, T1173o, H1141, H1144, H1140 and T1187o. For H1134, the MMscore is not significantly improved, but the interface as measured by the DockQ score [20] is. The suffices (o) from T1123o, T1173o and T1187o are excluded from the referrals throughout the rest of the text. H1129 was disregarded as it was out of memory using an NVIDIA A100 GPU with 40Gb of RAM, resulting in seven complexes in total. We used the same MSAs as AFsample (for consistency) and no template information. We only ran model_1 from AFM v2.0 and not all 5 models available from AFM. The MSAs from AFsample are available through: http://bioinfo.ifm.liu.se/casp15/. The MMscores for AFsample and AFM in Table 1 were obtained from https://predictioncenter.org/casp15.

The best models from CASP15 were used as references as these are all highly accurate and the real structures are not made available yet. Note that, alternative conformations may be present here, as the first and second ranked models from AFsample display completely different binding sites (e.g. H1144, Supplementary Figure 1) [8].

#### AlphaFold-multimer benchmark set with 2-6 chains

It is necessary to have an evaluation set of sufficient size for multimers. Therefore, we use a set from a recent benchmark of AlphaFold-multimer (AFM) of 2-6 chains that has been homology reduced on the structural level with pairwise MMscores<0.6. We use AFM v2 for comparison as v2.3 has a cutoff date of 2021-09-30 and v2 of 2018-04-30, leading to much fewer structures being available for evaluation with v3. In addition, the improved models in CASP15 from AFsample all came from AFM v2 [8,15]. We selected the examples which have a DockQ score <0.23 and an MMscore <0.75 as these are considered to be unsuccessful [3], to see if we can improve them. From the 1928 complexes in total, 728 were unsuccessful. From the 728, 334 (46%) could be predicted using an NVIDIA A100 GPU with 40 Gb RAM within an 8h time limit. For AFProfile, the models with the highest confidence were selected and scored in all analyses.

### AlphaFold-multimer v2

To predict the structures, we used AlphaFold-multimer (AFM)[7] v2 and the parameters from 2022.03.02 available from: https://storage.googleapis.com/alphafold/alphafold_params_2022-03-02.tar. We did not use any template information and turned on dropout to introduce noise everywhere except for in the structural module [8]. We only used one set of weights (model_1) from the five available [7] and used the default MSA generation pipeline (below).

By searching various databases with several genetic search programs, multiple sequence alignments (MSAs, four in total) are created. Jackhmmer from HMMER3 [21] is used to create three different MSAs by searching the databases Uniref90 v.2020_01 [22], Uniprot v.2021_04 [23] and MGnify v.2018_12 [24]. By searching the Big Fantastic Database [25] (BFD from https://bfd.mmseqs.com/) and uniclust30_2018_08 [26] together with HHBlits [27] (from hh-suite v.3.0-beta.3 version 14/07/2017) the fourth MSA is created. We used the reduced BFD option to save time and space. Species and positional information of genes is used to pair the results from the Uniprot search. The results from the other searches are block-diagonalized to maintain as much intra-chain information as possible. All four MSAs are used for the protein complex prediction by sampling and clustering the information.

### Finding the best profile residuals

The intuition behind directing the prediction process with a profile residual is that a random sample results in a structure that is accurate locally (i.e. for each single chain) but may not be so globally (the entire complex). The problem is to traverse the conformational landscape to the lowest valley, i.e. the best score [9,10]. The rationale we use is that AFM is capable of generating highly accurate predictions for all targets given informative sequences, but these have to be found. Another aspect is that the network needs to build up a sufficiently large internal representation of information. This is maintained by recycling and by resampling the MSA during each recycle. The MSA cluster profile bias is added in each recycle, which helps to direct the internal information to a more favourable direction.

One could sample diverse sequences randomly, but there are many possible combinations making the probability of finding the right ones unlikely. For this reason, many samples are drawn by AFsample. The most efficient way to direct the internal representation should be with gradient descent, using the fact that the network has learned the relationship between the MSA profile and the structure. One could traverse the MSA profile directly to generate a structure, but this may introduce issues between iterations due to different sequences being sampled. Therefore, we generate a bias to the MSA profile that learns to be robust across different samples and profiles.

We only use the first model (“model_1”) generated by AFM v2, although 5 models are available. The other four models are not explored to reduce computational cost and the fact that the weights should be highly similar and respond similarly to similar MSA profiles. We use a maximum of 100 iterations and Nvidia A100 GPUs with 40Gb RAM. The complexes that did not fit within these limitations were disregarded.

We explored varying the number of recycles between 0, 5, 10, 15 and 20 and using different learning rates (1e-4, 1e-3, 1e-2) with the Adam optimiser [28]. Note that only the gradients at the final pass through the network are used since we can’t take the gradients across recycles. This is consistent with the training procedure of AFM where only the gradients of the last recycle operation are used. The loss function used for all settings is confidence-1, where the confidence is defined as in equation 1.

It is not only important to obtain fast convergence, but also to obtain high scores. To analyse the relationship between confidence and model accuracy we used the MMscore from MMalign[2]. We use the best predictions from the CASP15 (Table 1) as the reference structures. The relationship between the MMscore and ranking confidence is noisy and in some cases it is decreasing (Supplementary Figure 2). For most targets, the MMscore increases with the ranking confidence (Supplementary Figure 3). The best Spearman correlation (0.65) is obtained from using a learning rate of 0.0001 and 0 recycles. However, a lr of 0.0001 and 20 recycles results in a higher density of highly accurate predictions at high confidences with a correlation of 0.59 which is why this is preferred (Figure 4).

**Figure 4.**
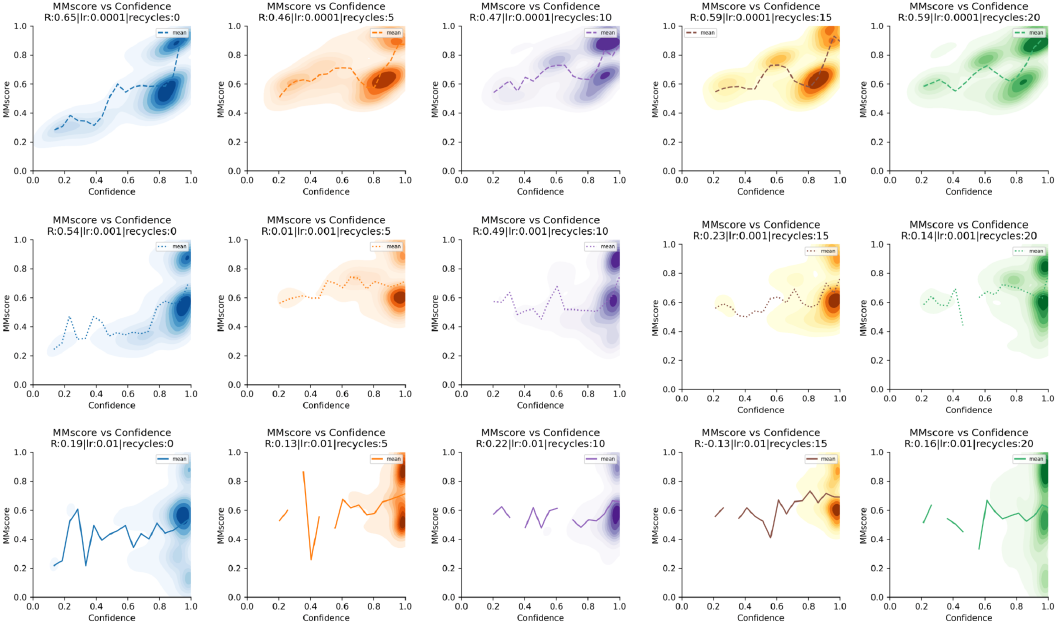
MMscore vs confidence for all 7 complexes using 100 iterations per complex (n=700) from the CASP15 set. The learning rate (lr) is increasing row-wise and the number of recycles column-wise. The Spearman correlations (R) are displayed in the figure legend. Some correlations are negative. The best correlation (0.68) is obtained from using lr=0.0001 and 0 recycles.

The correlations tend to decrease with an increasing learning rate, suggesting that it is not beneficial to traverse the loss landscape in too large steps, but rather to approach the highest confidence levels slowly. This is true for all targets except for H1141 where the optimisation procedure seems to get stuck at MMscores of 0.7 using a lr 0.0001 and a higher learning rate (0.01) results in MMscores>0.95.

In all analyses with AFProfile, the models with the highest confidence were selected and scored as the top models.

### Scoring

We used MMscore from MMalign[2] to score the predicted complexes. This score has a high correlation with the DockQ score[20] in the dataset we use for evaluation here (>0.8 for heteromers and >0.9 for homomers [3]). MMscore in addition captures alternative positions of identical chains (homomers), which DockQ does not, making it a better score for evaluation as such “swaps” should result in identical scores.

For the CASP15 structures, we can’t compare with the correct models as these are not available. However, the structures we selected for development (the highest scoring structures, Table 1) all report MMscores >0.9 and should therefore serve as adequate substitutes.

## Availability

The structures for the AFM benchmark with 2-6 chains are available from: https://gitlab.com/ElofssonLab/afm-benchmark

The code and instructions to run AFProfile are available from: https://github.com/patrickbryant1/AFProfile

The MMscores for AFsample and AFM in Table 1 and the models used as reference structures were obtained from: https://predictioncenter.org/casp15

The predicted models for the CASP15 and AFM benchmark set can be obtained from: https://zenodo.org/record/8108749

## Acknowledgements

This study was supported by the European Commission (ERC CoG 772230 “ScaleCell”), MATH+ excellence cluster (AA1-6, AA1-10), Deutsche Forschungsgemeinschaft (SFB 1114/C03). Computational resources were obtained from ZIH (SCADS) at TU Dresden with project id p_scads_protein_na.

## Contributions

P.B. designed and performed the studies, prepared all figures and wrote the initial draft of the manuscript which was later improved by all authors.

## Supplementary material

**Supplementary Figure 1.**
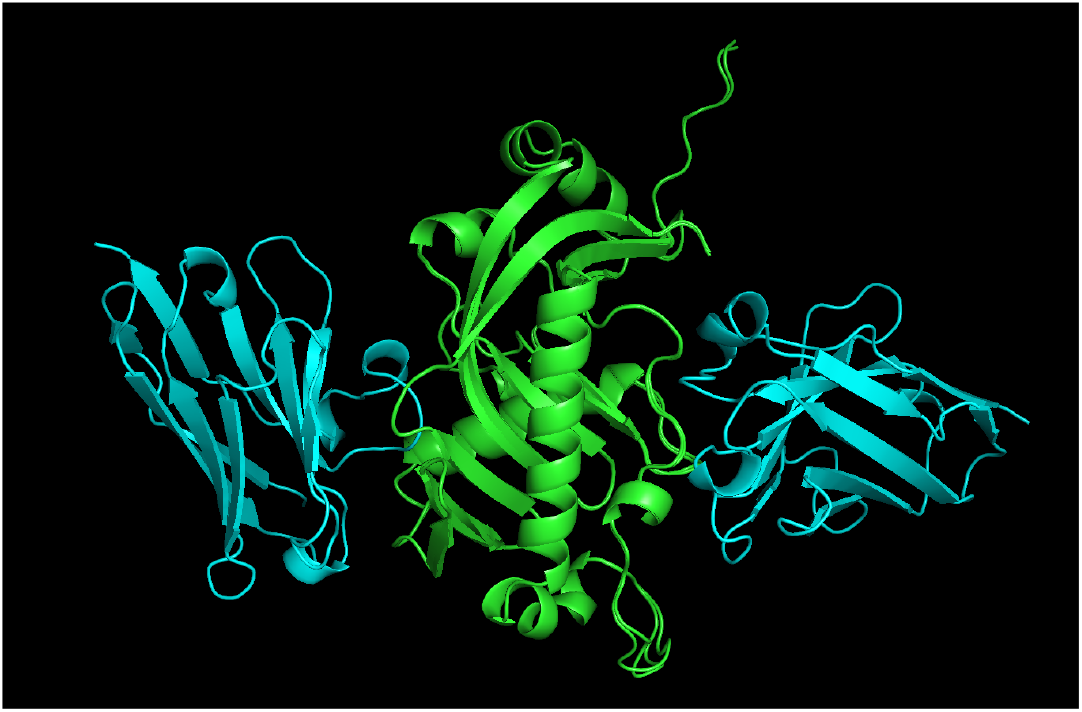
First (blue chain to the left) and second (blue chain to the right) ranked models superposed on the green chain from AFsample for target H1144.

**Supplementary Figure 2.**
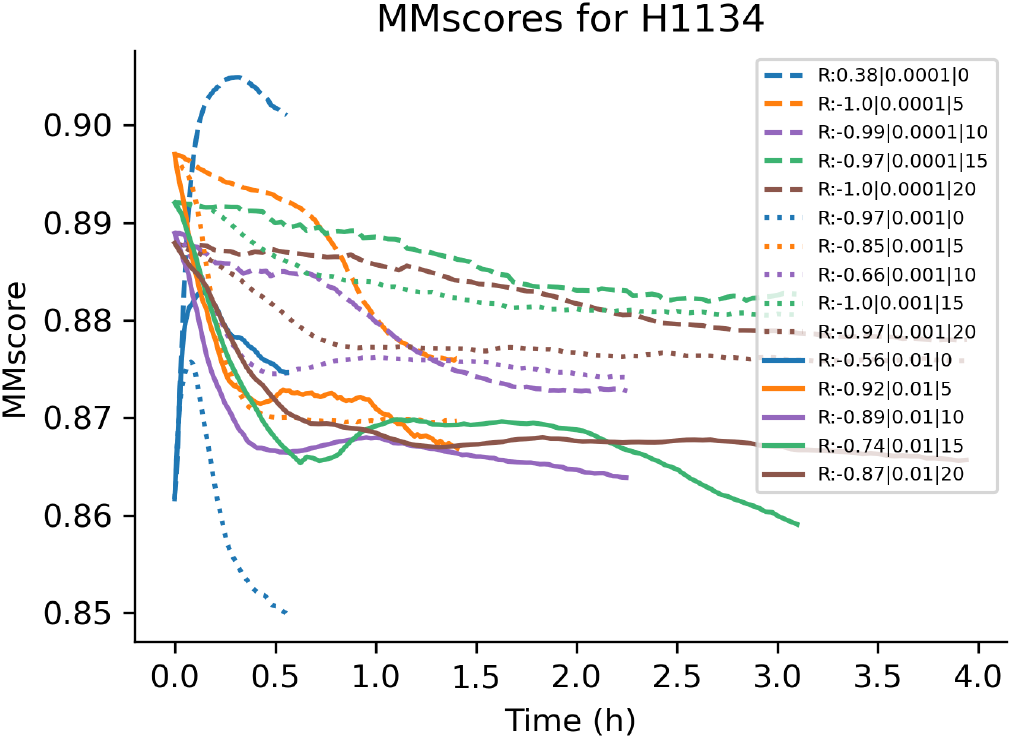
MMscore vs time for H1134 using different learning rates and recycles. The Spearman correlations (R) are displayed in the figure legend. The MMscore is stable when a good starting position is found for lower learning rates.

**Supplementary Figure 3.**
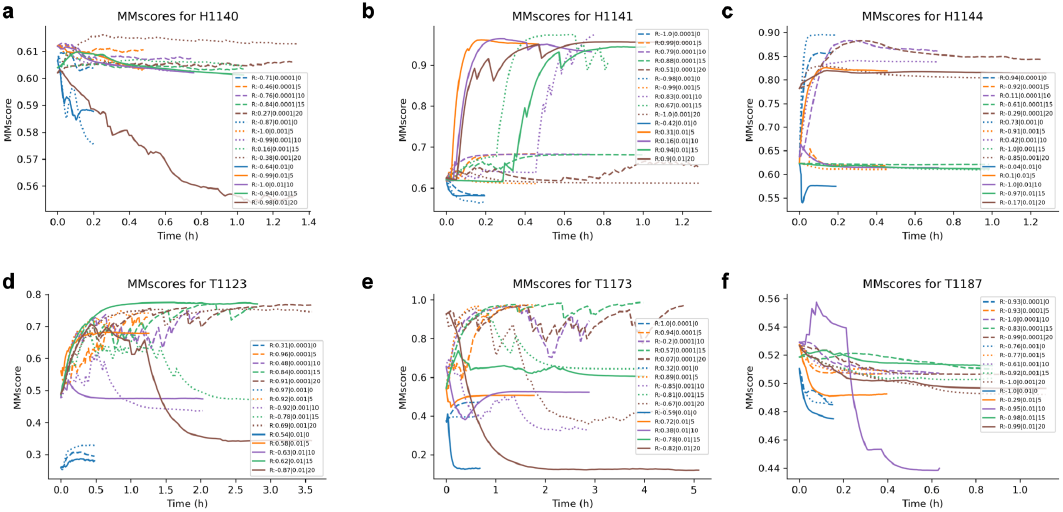
MMscore vs time for the targets H1141, H1144, H1140, T1123, T1173 and T1187 (a-e). The Spearman correlations (R) are displayed in the figure legend. The MMscore increases with time and thereby confidence for some settings in all cases except for T1187.

## Notes

### Competing Interest Statement

The authors have declared no competing interest.

## References

1. Jumper J, Evans R, Pritzel A, Green T, Figurnov M, Ronneberger O, et al. Highly accurate protein structure prediction with AlphaFold. Nature. 2021;596: 583–589.

2. Mukherjee S, Zhang Y. MM-align: a quick algorithm for aligning multiple-chain protein complex structures using iterative dynamic programming. Nucleic Acids Res. 2009;37: e83.

3. Zhu W, Shenoy A, Kundrotas P, Elofsson A. Evaluation of AlphaFold-Multimer prediction on multi-chain protein complexes. bioRxiv. 2022. p. 2022.12.08.519586. doi:10.1101/2022.12.08.519586

4. Bryant P, Pozzati G, Elofsson A. Improved prediction of protein-protein interactions using AlphaFold2. Nat Commun. 2022;13: 1–11.

5. Burke DF, Bryant P, Barrio-Hernandez I, Memon D, Pozzati G, Shenoy A, et al. Towards a structurally resolved human protein interaction network. Nat Struct Mol Biol. 2023;30: 216–225.

6. Akdel M, Pires DEV, Pardo EP, Jänes J, Zalevsky AO, Mészáros B, et al. A structural biology community assessment of AlphaFold2 applications. Nat Struct Mol Biol. 2022;29: 1056–1067.

7. Evans R, O’Neill M, Pritzel A, Antropova N, Senior A, Green T, et al. Protein complex prediction with AlphaFold-Multimer. bioRxiv. 2022. p. 2021.10.04.463034. doi:10.1101/2021.10.04.463034

8. Wallner B. AFsample: Improving Multimer Prediction with AlphaFold using Aggressive Sampling. bioRxiv. 2022. p. 2022.12.20.521205. doi:10.1101/2022.12.20.521205

9. Johansson-Åkhe I, Wallner B. Improving peptide-protein docking with AlphaFold-Multimer using forced sampling. Frontiers in bioinformatics. 2022;2. doi:10.3389/fbinf.2022.959160

10. Roney JP, Ovchinnikov S. State-of-the-Art Estimation of Protein Model Accuracy Using AlphaFold. Phys Rev Lett. 2022;129: 238101.

11. Frank C, Khoshouei A, de Stigter Y, Schiewitz D, Feng S, Ovchinnikov S, et al. Efficient and scalable de novo protein design using a relaxed sequence space. bioRxiv. 2023. p. 2023.02.24.529906. doi:10.1101/2023.02.24.529906

12. Bryant P, Elofsson A. EvoBind: in silico directed evolution of peptide binders with AlphaFold. bioRxiv. 2022. p. 2022.07.23.501214. doi:10.1101/2022.07.23.501214

13. Jendrusch M, Korbel JO, Kashif Sadiq S. AlphaDesign: A de novo protein design framework based on AlphaFold. bioRxiv. 2021. p. 2021.10.11.463937. doi:10.1101/2021.10.11.463937

14. Goverde CA, Wolf B, Khakzad H, Rosset S, Correia BE. De novo protein design by inversion of the AlphaFold structure prediction network. Protein Sci. 2023;32. doi:10.1002/pro.4653

15. Progress at protein structure prediction, as seen in CASP15. Curr Opin Struct Biol. 2023;80: 102594.

16. Zhang Y, Skolnick J. TM-align: a protein structure alignment algorithm based on the TM-score. Nucleic Acids Res. 2005;33: 2302–2309.

17. alphafold/docs/technical_note_v2.3.0.md at main · deepmind/alphafold. In: GitHub [Internet]. [cited 19 Jun 2023]. Available: https://github.com/deepmind/alphafold

18. Wayment-Steele HK, Ovchinnikov S, Colwell L, Kern D. Prediction of multiple conformational states by combining sequence clustering with AlphaFold2. bioRxiv. 2022. p. 2022.10.17.512570. doi:10.1101/2022.10.17.512570

19. del Alamo D, Sala D, Mchaourab HS, Meiler J. Sampling alternative conformational states of transporters and receptors with AlphaFold2. 2022 [cited 28 Jun 2023]. doi:10.7554/eLife.75751

20. Basu S, Wallner B. DockQ: A Quality Measure for Protein-Protein Docking Models. PLoS One. 2016;11: e0161879.

21. Eddy SR. Accelerated Profile HMM Searches. PLoS Computational Biology. 2011;7: e1002195.

22. Suzek BE, Huang H, McGarvey P, Mazumder R, Wu CH. UniRef: comprehensive and non-redundant UniProt reference clusters. Bioinformatics. 2007;23: 1282–1288.

23. UniProt Consortium. UniProt: the universal protein knowledgebase in 2021. Nucleic Acids Res. 2021;49: D480–D489.

24. Mitchell AL, Almeida A, Beracochea M, Boland M, Burgin J, Cochrane G, et al. MGnify: the microbiome analysis resource in 2020. Nucleic Acids Res. 2020;48: D570–D578.

25. Steinegger M, Mirdita M, Söding J. Protein-level assembly increases protein sequence recovery from metagenomic samples manyfold. Nature Methods. 2019;16: 603–606.

26. Mirdita M, von den Driesch L, Galiez C, Martin MJ, Söding J, Steinegger M. Uniclust databases of clustered and deeply annotated protein sequences and alignments. Nucleic Acids Res. 2017;45: D170–D176.

27. Steinegger M, Meier M, Mirdita M, Vöhringer H, Haunsberger SJ, Söding J. HH-suite3 for fast remote homology detection and deep protein annotation. BMC Bioinformatics. 2019;20: 473.

28. Kingma DP, Ba J. Adam: A Method for Stochastic Optimization. 2014. Available: http://arxiv.org/abs/1412.6980

